# SepIVA has a function in activating the mycobacterial divisome, which is inhibited during DNA damage

**DOI:** 10.64898/2026.01.24.701465

**Authors:** Anusuya Nepal, Cara C. Boutte

## Abstract

Bacterial cell division is a tightly regulated process that is carried out by a complex of proteins called the divisome, which is assembled in a defined sequential order. Upon assembly the complex is allosterically activated, which stimulates cell wall synthesis at the division site. Bacteria inhibit division during DNA damage by blocking either divisome assembly or activation. While the regulation of cell division is known to be important during *M. tuberculosis* infection, little is known about mycobacterial mechanisms of divisome function and regulation in DNA damage. By using *M. smegmatis* as a model organism, we find here that divisome factor SepIVA is involved in septum initiation, and that it is recruited to the mid-cell by FtsQ but it is not a recruitment factor itself. We find that a *sepIVA* loss-of-function defect can be suppressed by overexpression of *ftsW*, supporting a role for SepIVA in activation of the divisome complex. When cell division is inhibited during DNA damage, we find that SepIVA is delocalized from the division site, while the septal localization of FtsZ, FtsQ and FtsW are not impacted. We also find that the interaction between FtsQ and SepIVA is inhibited during DNA damage. Our results suggest that SepIVA is a trigger factor for activation of cell division during normal growth, and show that the signaling to inhibit cell division during DNA damage involves inhibition of its interaction with FtsQ.

**IMPORTANCE:** Cell division is critical for bacterial cells to propagate and cause infection. Despite its importance, division is a dangerous process as it requires building and subsequently hydrolyzing new cell wall material, in a place where the chromosome typically resides. If cell division is done before chromosome segregation is completed, or if cell wall metabolism is improperly regulated, then cell death results. The divisome is a protein complex that regulates cell division - coordinating it with the status of the chromosome and ensuring that the cell wall metabolic enzymes are carefully controlled. Many of the proteins in the divisome complex are highly conserved across bacterial Phyla; however, the factor that stimulates the complex to activate and initiate cell wall synthesis is not widely conserved. Here, we study the activation of the divisome in *Mycobacterium smegmatis*, a model for cell physiology of *Mycobacterium tuberculosis*. We find evidence that SepIVA, a protein found only in Actinobacteria, is likely the factor that stimulates activation of the divisome in mycobacteria. We also show that the association of SepIVA with the divisome is blocked under DNA damage, when cell division is inhibited. These results provide a model for the regulation of cell division in mycobacteria in growth and stress, and also provide insights into how bacteria with different types of septa may regulate division differently.

## INTRODUCTION

Cell division in bacteria is a highly controlled and regulated process. It is carried out by a multiprotein transenvelope complex called the divisome (1–4). The complex includes cytoskeletal factors, transmembrane proteins, and peptidoglycan synthases and hydrolases (5, 6) that work together to ensure that cell division occurs in the correct time and place, and that the integrity of the cell wall is maintained during the division process. Cell division is carefully regulated in response to nutrient and stress conditions (7–14)

Much of our understanding of bacterial cell division comes from *E.coli* - using it as a model we can divide essential divisome factors into functional categories. There are proteins that build the septum, and those that resolve the septum (15, 16). Of septal construction factors, we can separate out the enzymes FtsW and FtsI, which do the chemical work of building the peptidoglycan (17–20), from the regulators that control the activity of the enzymes. Among regulators of septal construction, there are three functional categories. First, membrane anchors such as ZipA and FtsA identify and establish the division site at the mid-cell (21–23). Second, recruitment and scaffolding factors promote assembly of the divisome. Cytoskeletal protein FtsZ is the central scaffolding factor, and it recruits the series of essential transmembrane proteins in a defined sequence: first FtsK, then FtsQ, FtsL and FtsB, followed by FtsW with FtsI and finally FtsN (24–27). Finally, the third functional category is allosteric activation of the FtsWI enzymes. FtsN’s association with the divisome triggers conformational changes in FtsA and FtsQLB that activate the peptidoglycan synthases (28–31).

Several components of the *E.coli* divisome are conserved in Mycobacteria (32, 33), including FtsZ, FtsQ, FtsL, FtsB, FtsK, and FtsWI (33). Initial recruitment of FtsZ to the division site is mediated by SepF instead of FtsA/ ZipA (34). The recruitment ordering of FtsZ, FtsL, FtsQLB and FtsWI appears to be similar to that in *E. coli* (33). The step of sensing completion of assembly of the divisome and triggering activation of FtsWI is undescribed in Mycobacteria, as there is no homolog of FtsN (35). Inter-divisome interactions may contribute to allosteric activation: Mycobacterial FtsZ directly interacts with FtsW (36) and with FtsQ (37)though the function of these interactions is not clear yet.

SepIVA is a protein broadly conserved across actinomycetales (33, 38); however, its functional role does not appear to be conserved. In *M. smegmatis*, SepIVA is essential for division and localizes to the septum and sub-polar regions (33, 39). The *sepIVA* homolog in *S.venezuelae* does not have a role in division (38). In *M.smegmatis*, SepIVA also interacts with conserved divisome protein FtsQ (33, 40). Depletion of SepIVA does not impact the formation of the FtsZ ring, indicating it is not a membrane anchor or early recruitment factor. SepIVA consists of a DivIVA-like domain and no predicted enzymatic activity(33). Its molecular function has not been described.

Cell division must occur in co-ordination with DNA replication for the next generation to receive an intact copy of the genetic material (41, 42). During DNA damage, cell division in bacteria is typically halted until the DNA is repaired (42). In rod shaped bacteria such as *E. coli*, *B. subtilis* and *C. crescentus*, this results in long and filamented cells, as elongation continues(43–46). In *E.coli*, cell division is inhibited when DNA damage activates expression of SulA (Huisman and D’Ari, 1981), which blocks division by inhibiting the assembly of FtsZ (47). In *C. crescentus*, two transmembrane proteins, SidA and DidA, work together to inhibit cell division through interaction with the division subcomplex FtsW/I/N (10, 48). In *B.subtilis*, SOS-induced YneA interacts with FtsL and PBPs to block divisome function (49–51). Thus, different organisms can inhibit different steps of the division process during DNA damage. In Mycobacteria ChiZ has been proposed as the checkpoint protein that inhibits cell division during DNA damage by destabilizing the FtsZ ring; however, a comprehensive model of its function are not described (52, 53).

In this work, we examined the assembly of the divisome under growth and DNA damage, with a focus on the role of SepIVA. We show that SepIVA is involved in initiation of septation, not septal resolution. Then, we examined the dependency of localization of several essential divisome factors and found that SepIVA is recruited by FtsQ to the mid-cell, but it is not itself a recruitment factor for essential divisome factors. We also showed that overexpression of FtsW could suppress a SepIVA loss-of-function mutant, which provides support for the model that SepIVA is an allosteric activator of FtsWI, likely through its interaction with FtsQ. We found that division is inhibited under DNA damage and that SepIVA is delocalized from the division site during DNA damage, while FtsZ, FtsQ and FtsW are not. We also found that the interaction between SepIVA and FtsQ is inhibited during DNA damage. Our results suggest that SepIVA is a trigger factor for activation of cell division during normal growth, and that its interaction with FtsQ is inhibited during DNA damage to inhibit cell division.

## RESULTS

### SepIVA is required for septal construction

SepIVA is an essential septation factor in mycobacteria (33), however its specific molecular function is poorly understood. To determine if SepIVA is required for septal cross-wall construction, or for septal resolution, we used an *Msmeg* strain in which SepIVA can be inducibly proteolyzed (33, 54).To visualize sites of active cell wall assembly following SepIVA depletion, we stained the cells with fluorescent D-amino acid HADA, which incorporates into nascent and remodeled peptidoglycan at the poles and septa (55, 56). Under SepIVA depletion, we observed a significant increase in the distance between the cell poles and the septum, as well as between successive septa, compared with control cells where SepIVA is at normal levels (Figure 1). The filamentation of the cells and the reduction in the HADA-labelled septa demonstrate that SepIVA is critical for the initiation of septal cross-wall formation, not for septal resolution.

**Figure 1.**
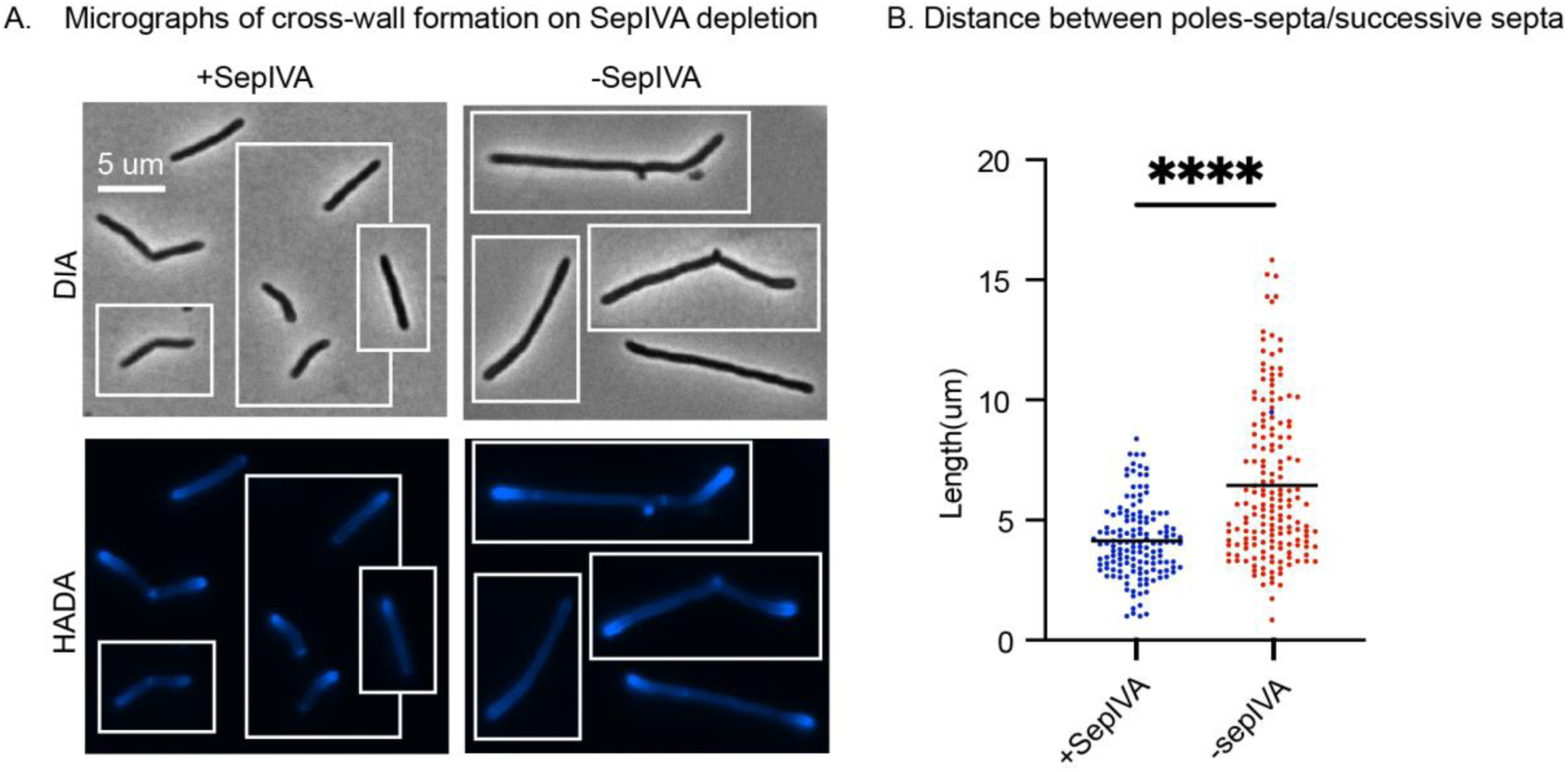
SepIVA is required for septal construction. **A.** Micrographs of the SepIVA-DAS strain with stable SepIVA-DAS (left) and depletion of SepIVA-DAS by SspB expression (right) stained with HADA. The scale bar is 5 micron and applies to all images. **B.** Distance between poles and septa or two successive septas of cells in A. Black bar represents the mean distance. 166 cells over two biological replicates of SepIVA-DAS depleted strains and 143 cells over two biological replicates of stable SepIVA-DAS strains were analyzed. ****,p<0.0001. p value was calculated by using unpaired t-test.

### SepIVA is recruited to the divisome site by FtsQ but does not recruit downstream divisome factors

In model species, essential non-enzyme septal construction proteins are involved either in 1) membrane anchoring of the Z-ring, 2) complex assembly (16, 57–59) or 3) allosteric activation of the cell division enzymes(28, 29, 60). We previously showed that SepIVA is not required for the septal localization of FtsZ, so it is not a membrane anchoring factor (33). Next, we tested if SepIVA is has a role in divisome assembly. We hypothesized that SepIVA would be recruited to the divisome adjacent to FtsQ, because it has been reported that SepIVA and FtsQ interact with each other (33, 40). To determine whether FtsQ recruits SepIVA in the divisome, we made an *Msmeg* strain, Ptet_OFF_:: *ftsQ* / mCherry-SepIVA, which allows transcriptional depletion of *ftsQ* and localization of SepIVA. In the control, mCherry-SepIVA is distributed mostly at the septal and sub-polar regions (33) (Fig. 2A), whereas depletion of *ftsQ* results in decreased localization of SepIVA at the midcell (Fig. 2 A,B). We conclude that SepIVA depends on FtsQ for its recruitment to the mid-cell, and is therefore a late recruit to the divisome.

**Figure 2.**
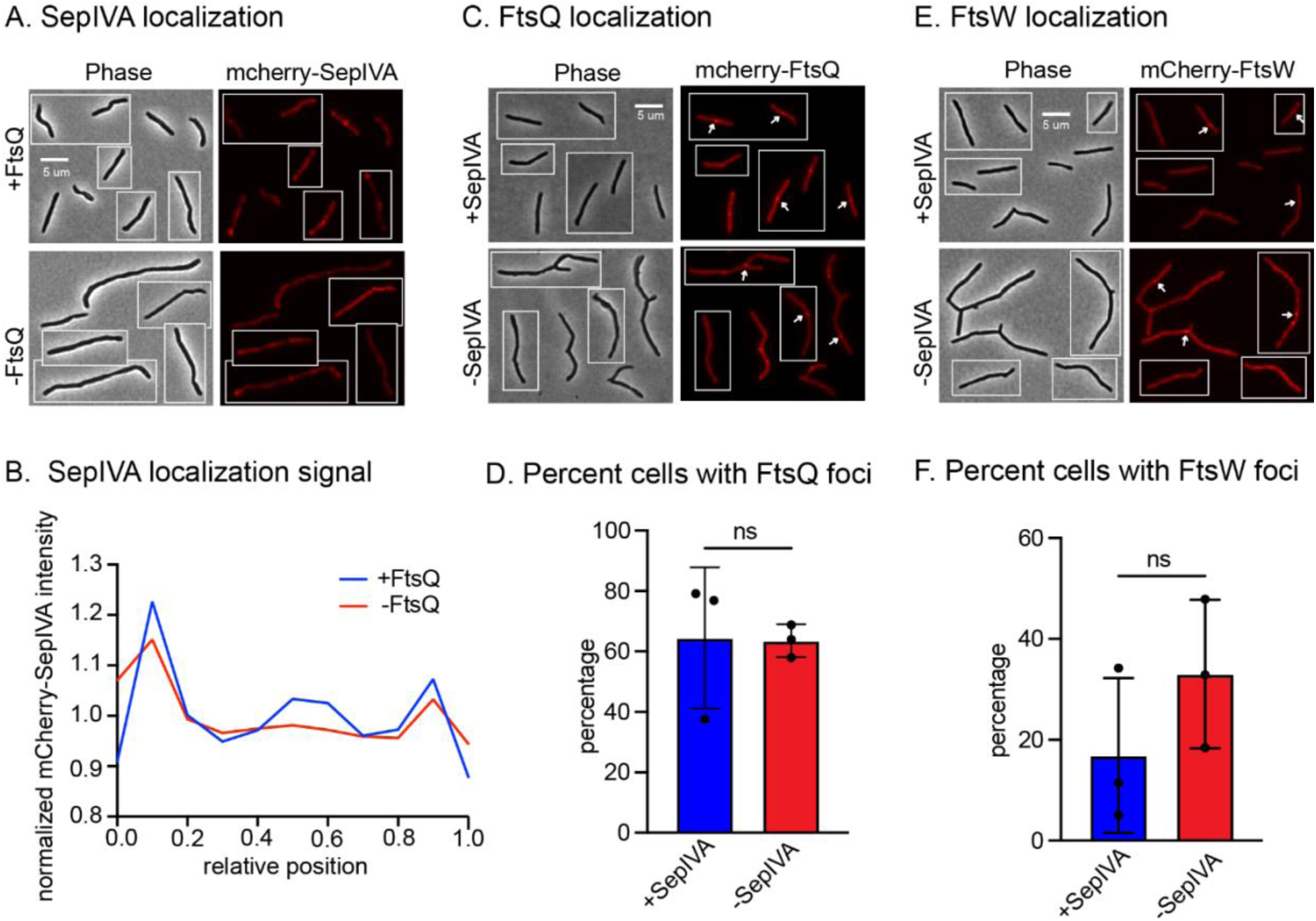
SepIVA is recruited to the divisome by FtsQ but does not recruit downstream divisome factors. **A.** Micrographs of Ptet_OFF_::*ftsQ* strains expressing mCherry-SepIVA in the induced (top, -ATC) and in transcriptional depletion of *ftsQ* (bottom, +ATC). Scale bar is 5 um and applies to all images in the figure. **B.** Mean Normalized intensity profile of mCherry-SepIVA signal along the relative position of ∼300 cells. Cells were sorted by poles such that the brighter HADA stained pole was set to 0 on X-axis. Normalized intensity values were calculated by dividing the mean intensity value of all cells at one position by the mean intensity value of all cells at all positions along the length of the cells. **C.** Micrographs of the SepIVA-DAS L5::mCherry-FtsQ strain with stable SepIVA-DAS (top) and SepIVA-DAS destabilized by SspB expression (bottom). The scale bar is 5 microns and applies to all images. Arrow heads point to mCherry-FtsQ foci. **D.** Percentage of total cells with FtsQ foci in SepIVA induced (blue) and SepIVA depleted (red) cells. 253 cells across three biological replicates of depleted and 274 cells across three biological replicates of controls strains were analyzed. ns, p>0.05 was calculated using an unpaired t-test. **E.** Micrographs of SepIVA-DAS L5::mCherry-FtsW strain with stable SepIVA-DAS (top) and SepIVA-DAS destabilized by SspB expression (bottom). The scale bar is 5 microns and applies to all images. Arrow heads point to mCherry-FtsW foci. **F.** Percentage of total cells with FtsW foci. 300 cells across three biological replicates of depleted and 240 cells across three biological replicates of controls strains were analyzed. ns, p>0.05 was calculated using an unpaired t-test.

Next we sought to determine if SepIVA has a role in recruitment or stabilization of FtsQ or FtsW to the divisome. To test this, we used *Msmeg* strains in which SepIVA-DAS could be degraded by SspB induction(33, 54). We then expressed either mCherry-FtsQ or mCherry-FtsW in this strain. We found that depletion of SepIVA did not affect the frequency of formation of FtsQ or FtsW foci (Figure 2 C,D,E,F). Our result that SepIVA does not stabilize FtsQ at the divisome corroborates earlier work from the Nandicoori lab (40) We also tried to check the localization of FtsI, the transpeptidase of the divisome, but we were unable to build a functional fusion construct of FtsI.

Mycobacterial FtsI has previously been shown to be recruited to the divisome by FtsW (61). The highly conserved nature of the FtsW-FtsI complex (20, 62) suggests that they are likely to behave in concert. Our data show that SepIVA is not a recruitment factor for FtsQ or FtsW, and we think it unlikely to be a recruitment factor for FtsI. By process of elimination, we therefore hypothesize that it is likely to be an allosteric activator of the divisome complex

### SepIVA‘s function in cell division is related to the activity of FtsW

To test if SepIVA is helping activate the FtsWI complex, we hypothesized that FtsW gain of function could bypass SepIVA’s loss of function, as has been observed in other species for *ftsN* loss of function (63, 64). We previously described the *sepIVA* R234M mutant, which is specifically partially defective in the cell division function of SepIVA (39). We used *Msmeg* strains expressing L5:: *sepIVA* WT or R234M with or without Ptet *ftsW*, to overexpress *ftsW* upon Atc treatment. We saw that the *sepIVA* R234M mutation resulted in cells that are longer than wild type, as we saw before (39). Overexpression of FtsW also resulted in cells that are longer than WT, which we expect is due to excess protein causing inhibitory misregulation of the complex. When we overexpressed FtsW in the *sepIVA* R234M mutant, the cell division defect was suppressed, resulting in cell lengths that are comparable to the WT (Fig. 3). Thus, overexpression of FtsW rescues SepIVA’ partial loss of function. We conclude from this result that SepIVA’s function is related to the activity of FtsW, and that it is likely to be involved in allosteric activation of the divisome complex.

**Figure 3.**
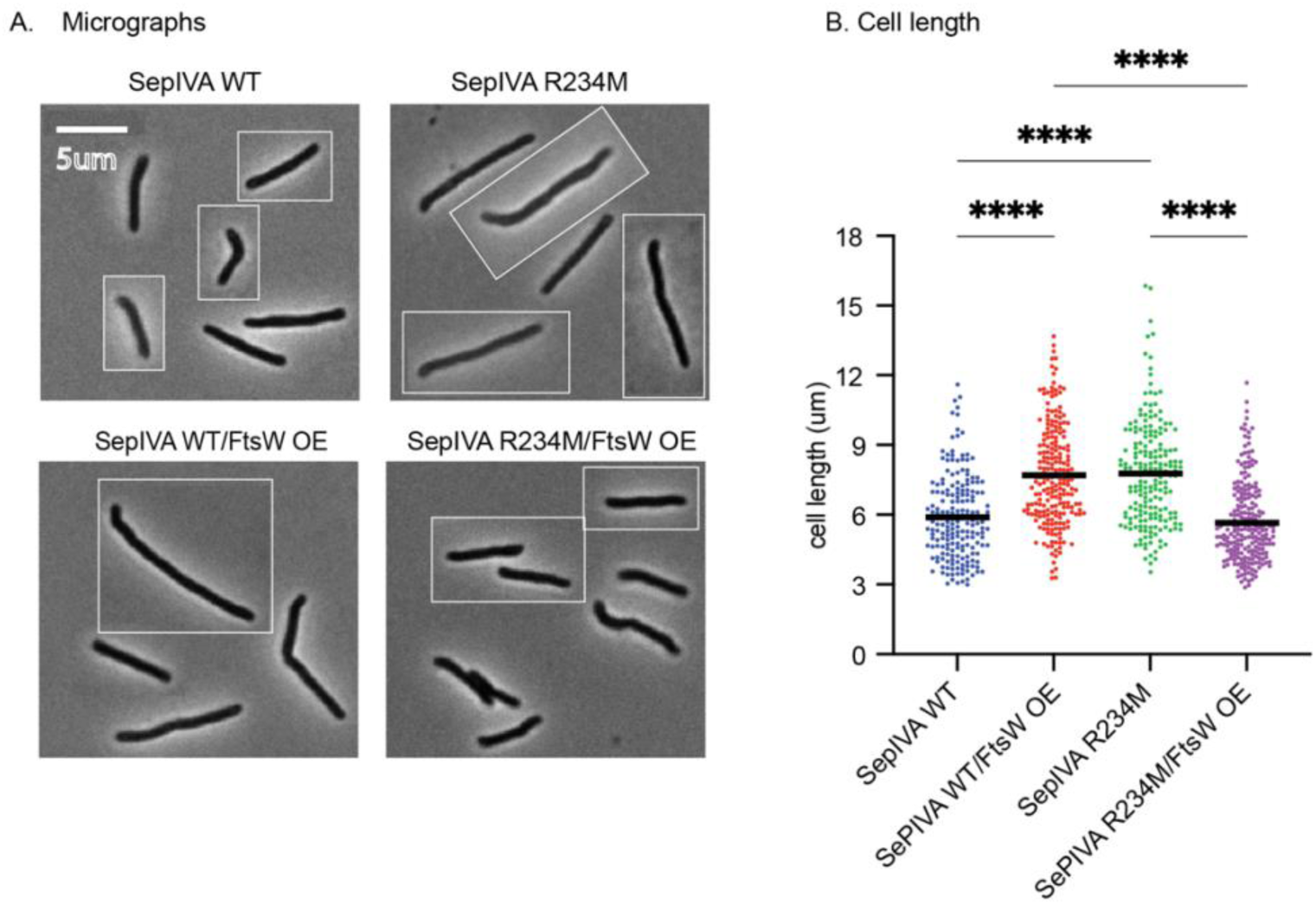
FtsW overexpression suppresses SepIVA’s cell division defects. **A.** Micrographs of *Msmeg* cells expressing L5:: *sepIVA* (WT or R234M) with or without Ptet FtsW, which causes FtsW overexpression upon anhydrotetracycline (Atc) treatment. All strains were treated with (Atc) for 16 hours. The scale bar of 5 micron applies to all images. **B**. Cell lengths of the strains from panel A. Bars represent the mean. 210 cells over three biological replicates for each strain were analyzed. ****, p<0.001 were calculated using an unpaired t-test.

### DNA damage does not impact septal localization of FtsZ, FtsQ or FtsW

DNA damage inhibits cell division in *Msmeg* as in many other species (42, 51, 63, 65) causing cells to elongate (52, 66) (Fig. 4). To determine whether DNA damage inhibits septal association of conserved divisome proteins, we used *Msmeg* strains expressing FtsZ-mCherry, GFP-FtsQ, or mCherry-FtsW as merodiploids. We grew the strains to log. phase and then treated with Mitomycin C (MitoC), which crosslinks DNA (67). For the control, the cells were left untreated. We found that FtsZ still retained its original septal position despite MitoC treatment (Figure 4a). Previous work suggested that ChiZ (Rv2719) inhibited cell division in DNA damage through inhibition of FtsZ assembly, though the effect of DNA damage alone on FtsZ assembly was never tested (52, 66). Our results suggest that ChiZ likely does not work through this mechanism, since FtsZ assembly is not inhibited in DNA damage. FtsQ foci were also present in most of the MitoC-treated cells. To determine if FtsQ foci were at the site of septation, we examined FtsQ co-localization with FtsZ, as FtsZ acts as a scaffold for FtsQ (33, 68). We saw that MitoC did not alter FtsQ-FtsZ co-localization (Fig 4c). Similarly, FtsW foci were still seen in the cells upon treatment with Mito C (Figure 4d), and there was no difference in their frequency between Mito C treated and control cells. (Figure 4e). These data show that septal localization of FtsZ, FtsQ or FtsW is not affected by DNA damage.

**Figure 4.**
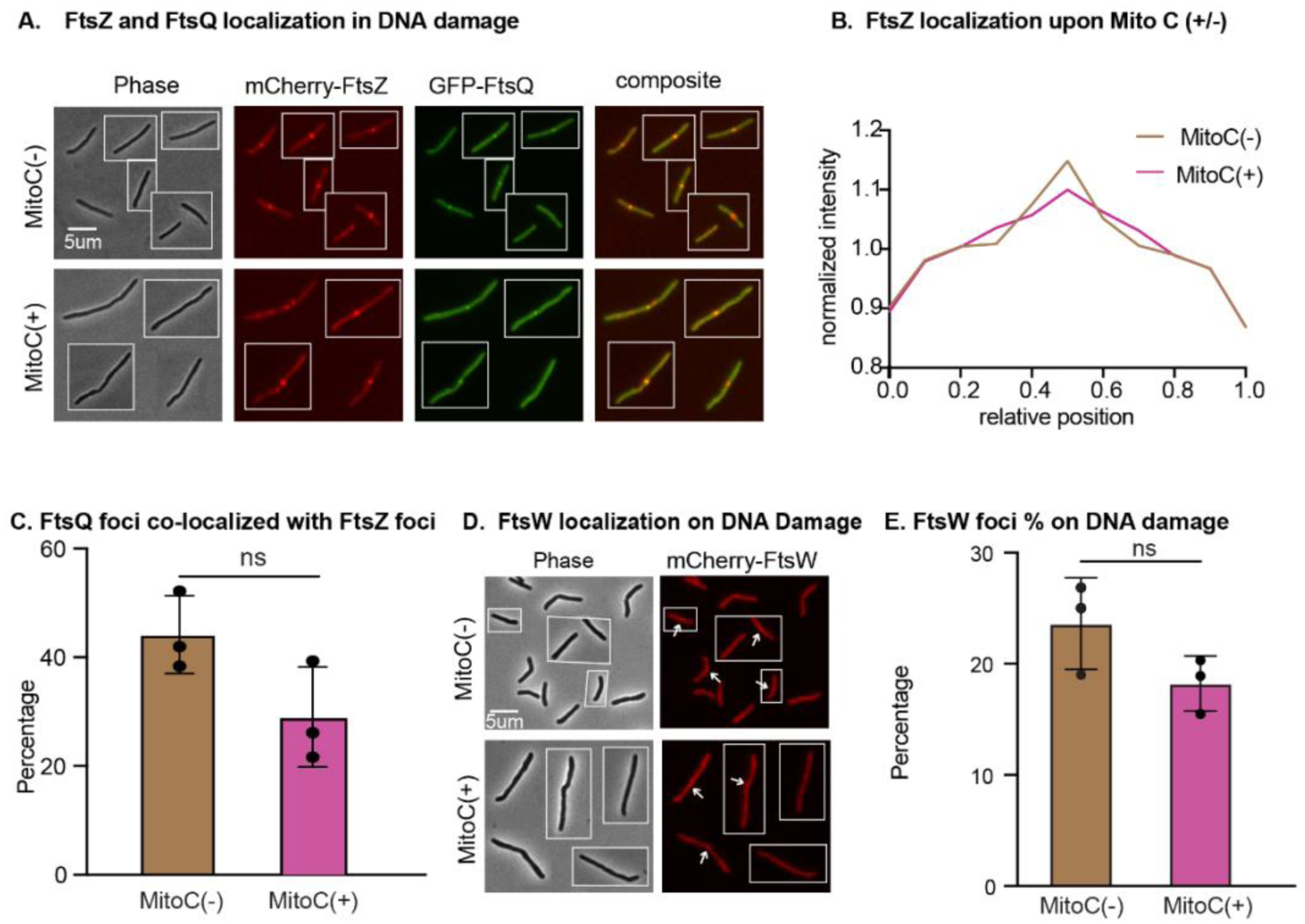
DNA damage does not impact septal localization of FtsZ, FtsQ and FtsW. **A.** Micrographs of *Msmeg* cells expressing FtsZ-mCherry and GFP-FtsQ. Scale bar is 5 microns and applies to all images. **B.** Normalized mean intensity profiles of FtsZ-mCherry signal along the length of 260 cells (untreated) or 284 cells (mitoC treated). Normalized intensity values were calculated by dividing the mean intensity value of all cells at one position by the mean intensity value of all cells at all positions along the length of the cells. **C.** The percentage of cells exhibiting FtsQ-FtsZ focal co-localization cells, among the cells with FtsZ foci. 358 cells across three biological replicates in MitoC untreated condition and 458 cells across three biological replicates in MitoC untreated conditions were analyzed. **D.** Percentage of mCherry-FtsW expressing cells that exhibit at least one FtsW focus under the two conditions. 240 cells across three biological replicates of MitoC untreated and 210 across three biological replicates cells of MitoC treated cells were analyzed. ns, p>0.05 was calculated using unpaired t-tests.

### DNA damage inhibits septal localization of SepIVA

Next, we sought to determine if DNA damage impacts the localization of SepIVA. We treated a strain expressing GFP-SepIVA with Mito C or UV. UV causes DNA damage by creating cyclobutene pyrimidine dimers (69). For the controls, the strains were left untreated. We find that SepIVA is delocalized from the mid-cell in both types of DNA damage (Fig:5). The quantified GFP-SepIVA intensity reveals reduced signal at the mid-cell for Mito C- and UV-treated cells, as compared to the pronounced peak in the untreated cells. (Figure 5b,d). Polar localization of SepIVA was not significantly altered. In conclusion, SepIVA is delocalized from the mid-cell in the presence of DNA damage.

**Figure 5.**
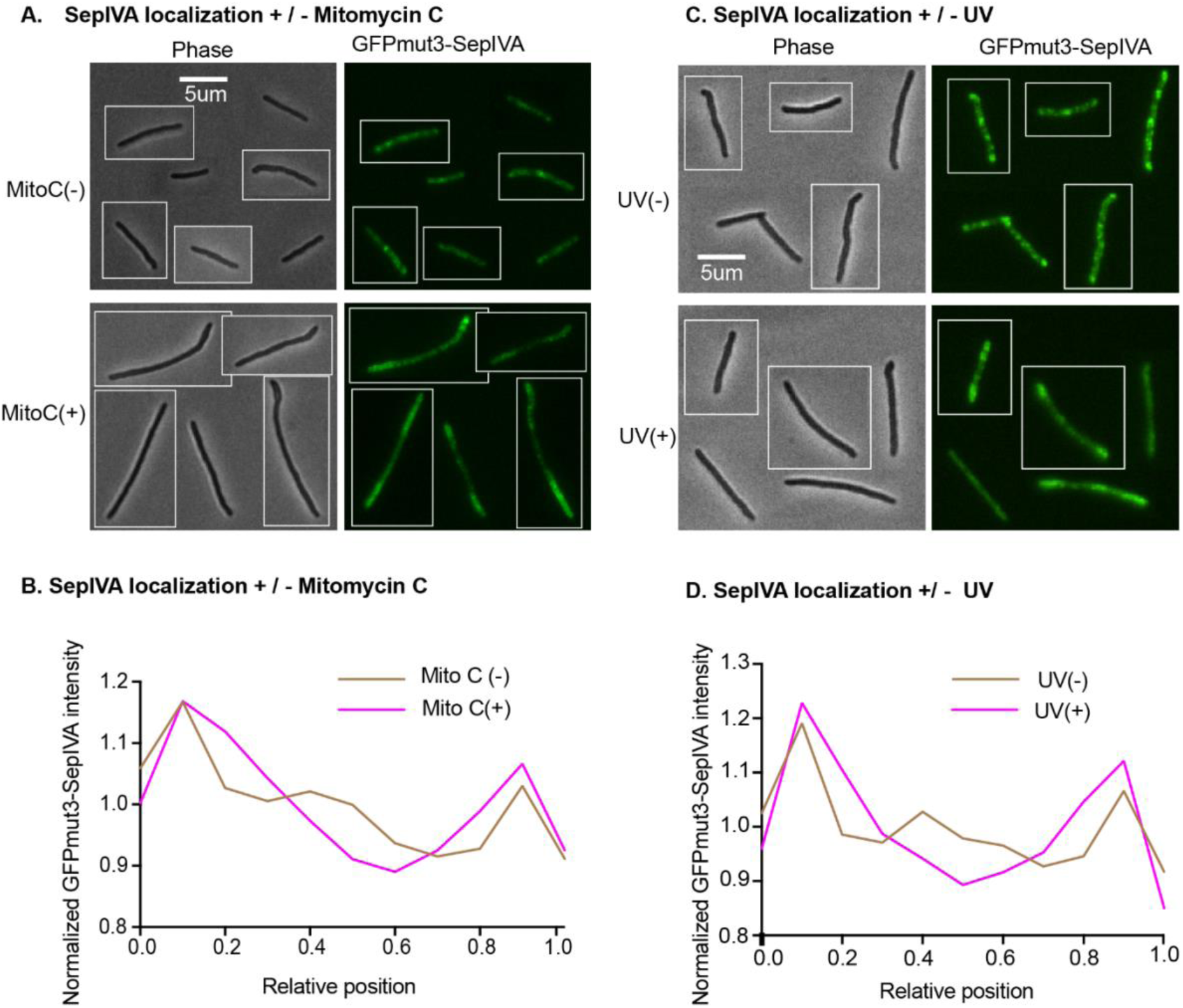
DNA damage inhibits septal localization of SepIVA. **A.** Micrographs of *Msmeg* cells expressing GFP-SepIVA before and after 4.5 hours of MitoC treatment at 350 ng/ mL. Scale bar is 5 microns and applies to all images. **B.** Normalized intensity profiles of GFP-SepIVA signal in untreated and mitoC-treated cells along the length of the cell, with the old pole set to 0 and the new pole set to 1 on the X axis. 310 cells for untreated and 340 cells for MitoC treated, across three biological replicates, were analyzed. Normalized intensity values were calculated by dividing each position’s intensity value by the mean intensity value of GFP-SepIVA signal across the cell. **C.** Micrographs of *Msmeg* cells expressing GFP-SepIVA before (top) and after (bottom) UV treatment of 150J for 10 seconds and outgrown for 3 hrs. Scale bar is 5 microns and applies to all images. **D.** Normalized intensity profiles of GFP-SepIVA signal before and after UV treatment, as in B. 284 cells for MitoC treated and 243 cells for untreated cells, across three biological replicates, were analyzed.

### DNA damage disrupts the interaction between SepIVA and FtsQ

Next we wanted to determine if SepIVA and FtsQ’s association is disrupted in DNA damage, given that SepIVA delocalizes from the mid-cell while FtsQ persists (Fig. 4ac, 5). We performed co-immunoprecipitations using an *Msmeg* strain expressing SepIVA-flag and FtsQ-strep as single copies, and a strain expressing only SepIVA-flag as a negative control. Under standard growth conditions, SepIVA-flag efficiently co-purified with FtsQ-strep (Figure 6), indicating a stable interaction between SepIVA and FtsQ even without using cross-linking as we did before (33). We then investigated whether the interaction is maintained during DNA damage. We observed that there was a significant reduction in the co-immunoprecipitation efficiency of SepIVA-Flag during UV-induced DNA damage compared to untreated controls (Figure 6). Since the stability of FtsQ-strep (bait protein) protein remained unaffected by UV treatment, it suggests that the observed decrease in SepIVA-FLAG recovery was due to a disruption of SepIVA-FtsQ interaction rather than a reduction in FtsQ level. These results demonstrate that while SepIVA and FtsQ form a strong complex during active growth, their interaction is inhibited under conditions of DNA damage. We propose that the inhibition of this interaction could result in inhibition of cell division because the divisome is not allosterically activated by association of SepIVA.

**Figure 6.**
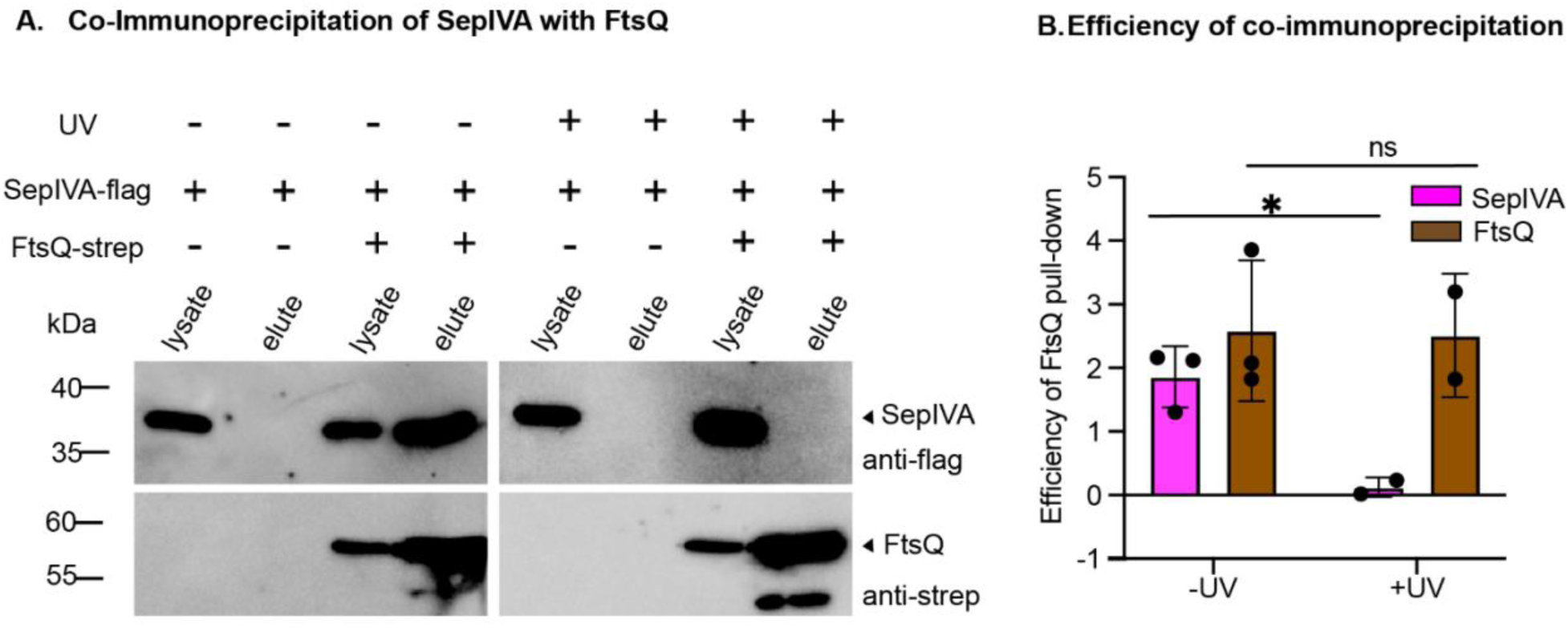
The interaction between FtsQ and SepIVA is inhibited during DNA Damage. A. α-FLAG and α-strep western blots of lysate controls and immunoprecipitation elutions from the SepIVA-flag (control) and SepIVA-flag/ FtsQ-strep (experimental) strains, using α-strep conjugated beads in the presence and absence of DNA damage (UV). The pulldown was performed twice, and a representative image is shown. B. The efficiency of the immunoprecipitation (FtsQ) or the co-immunoprecipitation (SepIVA), represented by the ratio of the elution peak band intensity to the lysate band intensity for each sample in A. Error bars represent standard deviation from two independent biological replicates. ns, non-significant (p<0.05); *, p<0.05 were calculated using an unpaired t-test.

## DISCUSSION

Our data support a model in which SepIVA’s interaction with FtsQ moves FtsQLB into an active conformation that activates the FtsIW complex to initiate peptidoglycan synthesis. First, SepIVA is a late-recruit to the divisome, like FtsN (25, 33, 70). It is recruited to the divisome complex by FtsQ but is not itself a recruitment factor to downstream factors like FtsQ or FtsW (Fig. 2). We were not able to test whether SepIVA recruits FtsI, because we were unable to make a functional fusion. However other work has shown that FtsI is recruited by FtsW (61, 62) and these proteins form a highly conserved protein complex which usually works together(20, 71). Though we do not expect that SepIVA is a recruitment factor for FtsI, we have not eliminated the possibility that there SepIVA could be recruiting FtsI or another undescribed divisome factor. The reduction in the HADA stained crosswalls (Fig. 1) and maintenance of mCherry-FtsW foci (Fig 2ef) in SepIVA depletion suggest that FtsIW are not actively polymerizing even if they are physically present in the divisome. We hypothesize that the interaction between SepIVA and FtsQ causes conformational rearrangements in the divisome complex that cue activation of the FtsWI enzymatic complex. The role of SepIVA in activation os supported by our data showing that in a *sepIVA* partial loss of function mutant (39), the diminished activation of FtsWI appears to be the rate-limiting step for septal peptidoglycan synthesis, since overexpression of FtsW in the *sepIVA* mutant strain suppresses the division defect (Fig 3).

In *E.coli*, the recruitment of FtsN to the divisome complex signals the completion of divisome assembly. Its association cues conformation changes in FtsA and the FtsQLB complex that ultimately activate the FtsWI enzymes (28, 60, 72–74). Mycobacteria do not have *ftsN* or a *ftsA* homolog. FtsA is important for stabilizing FtsZ in divisome complex (75), in addition to its role in allosteric activation (28, 60, 76). The role of FtsA in stabilizing the Z-ring is thought to be performed by SepF in Mycobacteria (34). How the Mycobacterial divisome is allosterically activated with neither FtsN nor FtsA has not been clear. However, other work has described direct interactions between FtsZ and FtsQ (37) and FtsZ and FtsW (61) which could help stabilize the FtsZ ring, or could be involved in allosteric activation of the complex. We propose that SepIVA may act as a trigger factor, inducing a conformational rearrangement of the divisome which could also involve these FtsZ-Q and FtsZ-W interactions.

FtsN is a transmembrane protein with a periplasmic peptidoglycan binding domain (77), while SepIVA is strictly cytoplasmic. Our model is that SepIVA’s triggering role in allosterically activating the divisome complex is simpler than FtsN’s role in both activating and coordinating septation. Through its peptidoglycan binding domain and transmembrane and cytoplasmic interactions with divisome factors, FtsN coordinates both septal synthesis and septal resolution, which occur nearly simultaneously in *E. coli* (78, 79). However, in mycobacteria the entire septum is built before it is cleaved (80) and therefore the mechanism to coordinate nearly-simultaneous septal construction and resolution is not required. Our model is that SepIVA triggers the active form of the divisome complex, but that it is not involved in coordinating septal resolution. This likely works similarly in *B. subtilis* and other species where the full septum is built before being resolved (81). Indeed, time lapse microscopy of divisome factors in *B. subtilis* indicates a simpler model for their motion dependent on septal peptidoglycan synthesis (82) than is seen in *E. coli* (79).

We found that the interaction between FtsQ and SepIVA, which we assume is key for triggering septation, was inhibited during DNA damage (Fig. 6). The localization data support this, showing that FtsQ is at the divisome without SepIVA during DNA damage (Fig. 4, 5). These results support the model that the interaction between FtsQ and SepIVA is a key regulatory event during Mycobacterial division. The association between FtsQ and SepIVA could be altered by post-translational modification, or by other proteins. SepIVA is extensively arginine methylated, and our previous studies indicate that these methylation sites regulate both elongation and division (39). FtsQ is phosphorylated, though its phosphorylation at threonine T24 (40, 83) does not affect its interaction with SepIVA (40); however, we have not yet tested whether DNA damage might activate other post-translational modifications. It might also be that a protein induced during DNA damage (84, 85) could block the interaction between FtsQ and SepIVA, analogous to similar inhibitory systems in *C. crescentus* (10) and *B. subtilis* (13). ChiZ is one protein that has been identified as a potential regulator of cell division in DNA damage (52, 53). It is upregulated during DNA damage stress and interacts with FtsQ (53); thus, one model could be that ChiZ inhibits the interaction between SepIVA and FtsQ. However, we found that DNA damage still substantially inhibits cell division in an *Msmeg* Δ*chiZ* strain (Fig. S3). Thus, if ChiZ does inhibit this interaction, it is not the only regulator involved.

## MATERIALS AND METHODS

### Bacterial strain and growth conditions

All *Msmeg* strains were cultured in 7H9 (Becton, Dickinson, Franklin Lakes, NJ) medium supplemented with 5 g/liter albumin, 2 g/liter glucose, 0.85 g/liter NaCl, 0.003 g/liter catalase, 0.2% glycerol, and 0.05% Tween 80. LB Lennox plates were used for plating *Msmeg* and *E. coli* strains. Genetically unstable *Msmeg* strains, such as the *sepIVA* R234M strain, were cultured directly from plates immediately after construction. *E.coli* strains, Top10, XL1-Blue, and DH5α, were used for cloning. Antibiotic concentrations used for mycobacterial selection were kanamycin, 25 μg/mL; hygromycin, 50 μg/mL; nourseothricin, 20 μg/mL; and zeocin, 20 μg/mL. For *E. coli*, kanamycin, 50 μg/mL; nourseothricin, 40 μg/mL; and zeocin, 25 μg/mL were used. Anhydrotetracyline (aTc) was used at between 50 ad 500 ng/ml for gene induction or repression.

### Strain construction

Plasmids were all made using Gibson cloning (86) (87). Vectors were sequence confirmed and transformed into electro-competent *Msmeg* strains. All strains used are listed in Table S1, plasmids used are listed in Table Y, and Primers used are listed in Table S3. The SepIVA depletion strain was previously described (33). The strain with a knockout of *ftsQ* and a complementation of *ftsQ* at the L5 site was also previously described (33). We used L5 allele swapping to exchange plasmids at the L5 site of this strain (88). We used ORBIT recombineering (89) to integrate a vector that attaches a FLAG-DAS tag at the C-terminus of *sepIVA* in a strain that expresses FtsQ-strep from the L5 site. The ORBIT primer is in Table Z. The Δ*chiZ* strain was built using double stranded recombineering (90)

### Cell staining

Cultures were grown to logarithmic phase and stained with 10 µM HADA (Tocris Bioscience) for 15 min with rolling and incubation at 37°C. HADA was then washed out, and the pellets were resuspended in 7H9 media. The cells were mounted on HDB agar pads before imaging. Images were either taken live or within 24 hrs of fixing them. Fixing of cells was performed by resuspending the pellets in PBS-Tween 80 with 1.6% paraformaldehyde solution for 15-20 minutes, followed by washing and resuspending them in PBS-Tween 80. The images were taken within 24 hrs of fixing.

### Microscopy and data analysis

Cells were immobilized on pads with 1.5% agarose in either HdB (for live-imaged samples) or PBS (for fixed samples) medium, and the images were taken on a Nikon Ti-2 widefield epifluorescence microscope with a Photometrics Prime 95B camera and a Plan Apo 100x, 1.45 Numerical Aperture (NA) lens objective. GFPmut3 fusions were imaged with a filter cube with a 470/40 nm excitation filter, a 525/50 nm emission filter, and a 495 nm dichroic mirror. mCherry2B fusionsions were detected using a filter cube with a 560/40 nm excitation filter, a 630/70 nm emission filter, and a 585 nm dichroic mirror. HADA was detected using a filter cube with a 350/50 nm excitation filter, a 460/50 nm emission filter, and a 400 nm dichroic mirror. All images were processed using NIS Elements software and analyzed using FIJI and MicrobeJ (Ducret et al., 2016) to create cell ROIs and extract fluorescence data from them. Mean intensities, maximum intensities, and minimum intensities of at least 250 cells per genotype were quantified using MicrobeJ.

Mean intensity profiles were plotted using the “XStatProfile” plotting tool in MicrobeJ. P-values were calculated using ordinary one-way ANOVA, Dunnett’s multiple comparisons test, with a single pooled variance using GraphPad Prism (v9.2). The mCherry-FtsQ foci were identified as foci that exceeded a threshold metric in which the ratio of the intensity of a focus over the mean intensity of a cell was greater than 1.65. The mCherry-FtsW foci in SepIVA depletion were identified as foci that exceeded a threshold metric in which the intensity of a focus over the mean intensity of a cell was greater than 1.55. The mCherry-FtsW foci in DNA damage were identified using a threshold metric in which the signal of a foci over the mean signal of a cell was greater than 1.44.

### Dependency of Localization Assays

For depletion of SepIVA and localization of FtsQ, 500ng/ml Anhydotetracycline (Atc) was added to the SepIVA depletion strain ( SepIVA-DAS) (CB 3290) in the early log. phase and incubated by rolling at 37°C for 7-8 hrs, stained with HADA for 15 minutes followed by washing of HADA and fixing the cells. Localization of FtsW in the same background was performed similarly as FtsQ except that the cells were fixed for half an hour. The images were taken within 24 hrs of fixing. Similarly, For FtsQ depletion in *Msmeg* Δ*ftsQ* PtetOFF:: FtsQ (CB 3537), the cells were grown to log. phase and treated with 500ng/ml Atc for 7-8 hrs to deplete FtsQ transcriptionally, HADA stained for 15 minutes followed by washing with 7H9 media. The images were taken immediately after mounting, with no fixation.

### DNA damage assays

For the DNA damage assays, strains were treated with 350ng/ml of Mitomycin C in log phase and rolled for 5 hours at 37°C. At 4 hours and 45 minutes after mitoC treatment, 10 µM HADA was added, and the cells were rolled for an additional 15 minutes, before being washed in 7H9 media and mounted for imaging. For UV treatment of the cells, the strains were first grown to OD 0.3, poured into a clean petri dish, exposed to UV in the UV crosslinker box (VWR) for 10 seconds at 150 J energy, then pipetted back into a culture tube. The culture was then incubated for 3 hours by rolling at 37°C before imaging. For control, the cells were not exposed to UV.

### Co-Immunoprecipitation

To test the interaction of SepIVA and FtsQ in normal and DNA-damaged conditions, 400ml culture of the ftsQ-strep SepIVA-flag strain (CB 3785) and SepIVA-flag strain (CB 1160) strains were grown to late log phase (OD6000.6-0.9). For the UV treated cultures, one half of each culture was poured into 245 mm × 245 mm petri plates and exposed to 150 J of UV light for 10 seconds. The cultures were then pipetted back into and flask and incubated for 1 hr 40 minutes with shaking at 37°C. Untreated cultures continued shaking at 37°C. The cultures were spun down and resuspended in Wash buffer (100 mM Tris pH 8.0,150 mM NaCl, 1 mM EDTA), mixed with protease inhibitor (Roche tablets and 1mM PMSF), and 0.1% n-dodecyl-β-D-maltose (Cayman Chemical). The cells were lysed by being passed twice through a French press, and the lysate was centrifuged at 15,000 rpm for 15 min at 4°C. The protein concentration in the supernatant was normalize to 60 mg of total protein for all samples, which were incubated with 200ul of Streptactin-covered beads (IBA) for 2 hrs in the cold room. The beads were washed twice with Wash buffer, and the protein was eluted with 40 ul of Buffer BX (IBA). Elutions were unfolded in Laemmli buffer and incubated at 95°C for 10 minutes before 15 ul of eluted samples were loaded onto an SDS gel. 30 ng of total protein in lysate controls were also loaded. After SDS-PAGE, proteins were transferred onto polyvinylidene difluoride (PVDF) membranes. The membranes were probed with anti-flag (Sigma, F1804) antibody (1:10,000 dilution in TBS pH 8.0) and anti-strep (Invitrogen, MA5-44988) antibody (1:10,000 dilution in TTBS), which were then probed with anti-mouse secondary antibody (Invitrogen, 31430).

